# Functional reconstitution of TatB into thylakoidal Tat translocase

**DOI:** 10.1101/708586

**Authors:** Sarah Zinecker, Mario Jakob, Ralf Bernd Klösgen

## Abstract

We have established an experimental system for the functional analysis of thylakoidal TatB, a component of the membrane-integral TatBC receptor complex of the thylakoidal Twin-arginine protein transport (Tat^1^) machinery. For this purpose, the intrinsic TatB activity of isolated pea thylakoids was inhibited by affinity-purified antibodies and substituted by supplementing the assays with TatB protein either obtained by *in vitro* translation or purified after heterologous expression in *E. coli*. Tat transport activity of such reconstituted thylakoids, which was analyzed with the authentic Tat substrate pOEC16, reached routinely 20 - 25% of the activity of mock-treated thylakoid vesicles analysed in parallel. In contrast, supplementation of the assays with the purified antigen comprising all but the N-terminal transmembrane helix of thylakoidal TatB did not result in Tat transport reconstitution which confirms that transport relies strictly on the activity of the TatB protein added and is not due to restoration of the intrinsic TatB activity by antibody release. Unexpectedly, even a mutant TatB protein (TatB,E10C) assumed to be incapable of assembling into the TatBC receptor complex showed low but considerable transport reconstitution underlining the sensitivity of the approach and its suitability for further functional mutant analyses. Finally, quantification of TatB demand suggests that TatA and TatB are required in approximately equimolar amounts to achieve Tat-dependent thylakoid transport.

## Introduction

Among the mechanisms catalyzing the transport of proteins across cellular membranes the Twin-arginine translocation (Tat) mechanism is distinguished by the fact that it can translocate proteins in a fully folded conformation (Clark and Theg, 1997; Hynds et al., 1998). It is found operating at the cytoplasmic membranes of bacteria and archaea as well as at the thylakoid membranes of chloroplasts and their progenitors, cyanobacteria (Müller and Klösgen, 2005). Despite some differences in detail, the overall structure of the Tat machinery as well as the presumed mechanism of membrane translocation is largely conserved among all these systems (e.g., Cline et al., 2015; Hauer et al., 2017; Petterson et al., 2018).

Tat machinery consists of three membrane integral subunits named TatA, TatB, and TatC, which in the thylakoid system are also addressed as Tha4, Hcf106, and cpTatC, respectively (Müller and Klösgen, 2005). TatC is a polytopic membrane protein of approximately 33 kDa with six transmembrane helices and a small N-terminal hydrophilic domain that is exposed to the cytosol of bacteria and the stroma of chloroplasts (Rollauer et al., 2012; Ramasamy et al., 2013). In contrast, TatA and TatB have each a single N-terminal transmembrane helix followed by a short hinge region, an amphipathic helix, and a largely unstructured C-terminal hydrophilic domain that is likewise situated on the cytosolic/stromal side of the membrane (Hu et al., 2010; Rodriguez et al., 2013; Zhang et al., 2014; Pettersson et al., 2018). Together with the considerable sequence homology in the helical domains, this common structure indicates that the genes of TatA and TatB are probably derived from a common ancestor (Yen et al., 2002; Alcock et al., 2016).

In spite of this close relationship, TatA and TatB fulfil distinct functions in the transport process. TatA is responsible in a yet unknown manner for the actual membrane translocation step. Three models of TatA activity have been proposed, namely (i) TatA constituting an adaptable translocation pore that is temporarily formed on demand (Mori and Cline. 2002; Gohlke et al., 2005; Cline and McCaffery, 2007), (ii) local weakening of the membrane at the sites of TatA accumulation which allows protein translocation directly through the lipid bilayer (Natale et al., 2008; Hou et al., 2018), or (iii) TatA providing the (co-)enzymatic activity converting the entire Tat apparatus into the active translocase (Jakob et al., 2009; Hauer et al., 2013; 2017). The latter is supported by the observation that TatA is not strictly associated with the thylakoid membrane but found in active form also soluble in the chloroplast stroma (Frielingsdorf et al., 2008). TatB, on the other hand, is always membrane-bound in all systems characterized to date. Together with TatC it constitutes heterooligomeric integral membrane complexes of approximately 560 - 700 kDa which provide the receptor structures recognized by the Tat substrates (Berghöfer and Klösgen, 1999; Cline and Mori, 2001). In spite of these differences, both, TatA and TatB, were recently shown by cross-linking experiments in *E. coli* to interact with TatC and it was suggested that they might swap their positions at TatC during the transition of the translocase from the resting state into the active state (Habersetzer et al., 2017).

In the thylakoid system, analysis of Tat transport rests mainly on transport experiments performed with isolated thylakoid vesicles and Tat substrates generated by *in vitro* transcription/translation. These *in thylakoido* assays were shown to facilitate almost quantitative membrane transport of a large number of authentic and chimeric Tat substrates (e.g., Cline et al., 1992; Klösgen et al., 1992; Fan et al., 2008). They were furthermore shown to be suitable also for the functional analysis of one of the Tat subunits, namely TatA. In so-called *in thylakoido* reconstitution assays, the intrinsic TatA activity of thylakoid membranes was eliminated by extraction with solutions of chaotropic salts or high pH (Frielingsdorf et al., 2008), or by treatment with specific anti-TatA antibodies (Dabney-Smith et al., 2003; Hauer et al., 2013). Subsequently, transport was reconstituted by supplementing the assays with soluble TatA obtained from *in vitro* translation or heterologous overexpression. The antibody approach turned out to be particularly suitable and was used, for example, for the characterization of mutant or chimeric TatA derivatives (Dabney-Smith et al., 2003; Hauer et al., 2017) as well as for the quantification of TatA demand during transport of a model Tat substrate (Hauer et al., 2013).

Here we have adapted this approach to the analysis of TatB function. Like with TatA, intrinsic thylakoidal TatB activity was inhibited by specific anti-TatB antibodies, which prevents protein transport by the Tat pathway. Reconstitution of Tat-dependent membrane transport was subsequently achieved by supplementing the assays with soluble TatB proteins facilitating the analysis of the demand and catalytic activity of this Tat subunit during transport of a model Tat substrate.

## Results

### Thylakoidal TatB can be functionally replaced by externally added protein

Reconstitution experiments, which are suitable for the functional analysis of single subunits of a protein transport machinery, depend on an assay system that is devoid of the intrinsic activity of the subunit under study as well as on the availability of this protein in sufficient amounts and activity to allow for transport recovery. In the case of thylakoidal TatA, this was achieved by inhibiting the intrinsic TatA activity by specific antibodies and reconstituting the transport properties of these thylakoid vesicles by the addition of TatA protein obtained from *in vitro* synthesis or bacterial overexpression (Dabney-Smith et al., 2003; Hauer et al., 2013; 2017).

To allow for a similar analysis also of thylakoidal TatB, specific antibodies had to be raised. For this purpose, we have overexpressed in *E. coli* a C-terminally His-tagged derivative of pea TatB comprising all but the N-terminal transmembrane helix (TMH) of the mature protein (Fig. 1A, antigen). The resulting polypeptide of approximately 18 kDa was purified by a combination of affinity chromatography, preparative SDS-PAGE, and reversed phase HPLC, in analogy to the procedure described for the antigen of TatB from *Arabidopsis thaliana* (Jakob et al., 2009). The purified protein was then used for immunization of rabbits as well as for the subsequent affinity purification of the TatB-specific antibodies from the antisera.

**Fig. 1.**
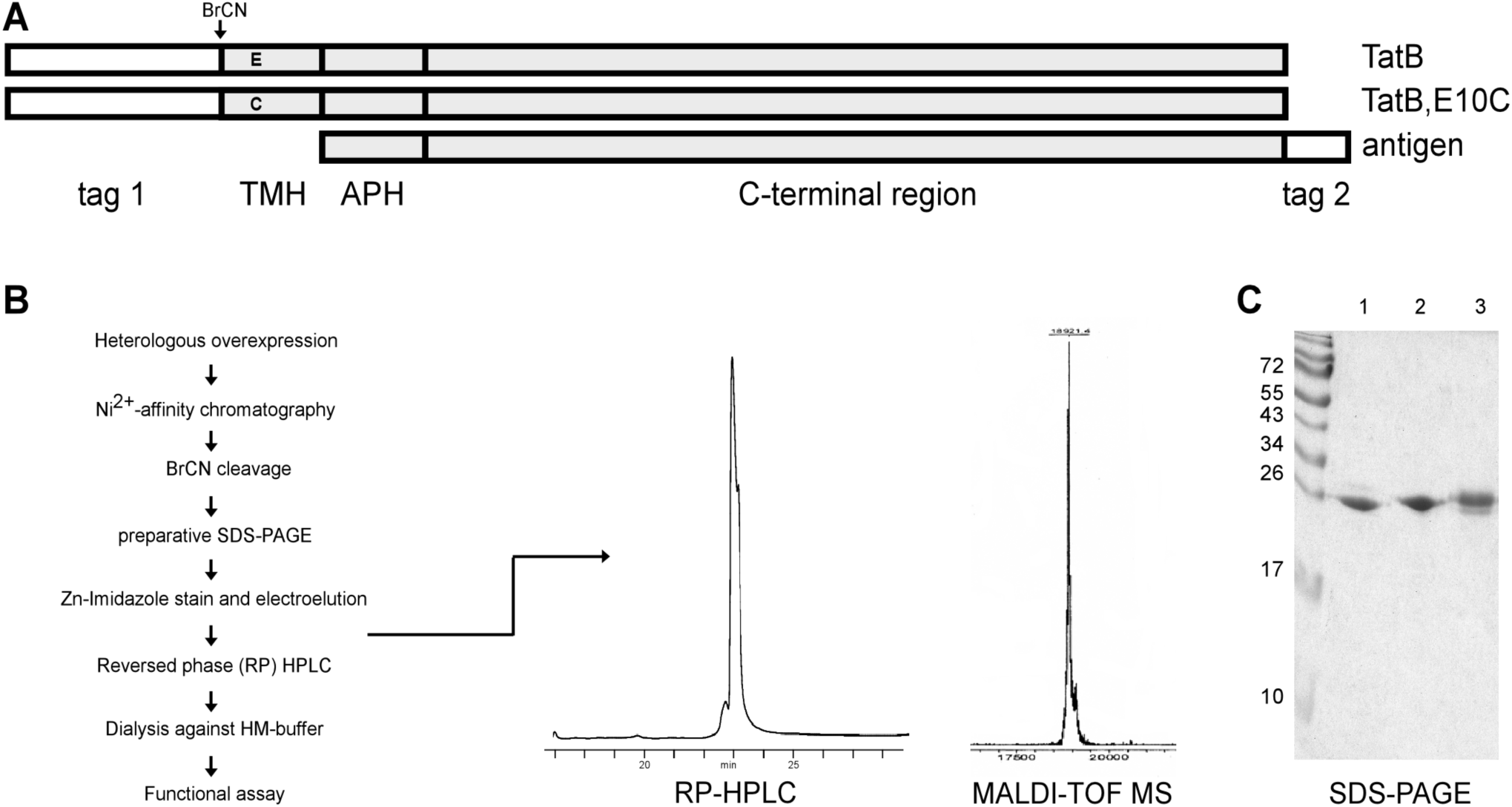
Heterologous overexpression and purification of TatB and TatB derivatives from pea. (**A**) Schematic representation of the constructs encoding *TatB, TatB,E10C*, and TatB *antigen* used for bacterial overexpression. The two full-size proteins comprising the N-terminal transmembrane helix (*TMH*), the amphipathic helix (*APH*), and the hydrophilic *C-terminal region* were expressed with an N-terminal His-/S-peptide-tag (*tag 1*) which can be removed by BrCN cleavage (*BrCN*) to yield the respective mature protein. The TatB antigen, which lacks the N-terminal TMH, carries a non-cleavable His-tag at its C-terminus (*tag 2*). (**B**) Flow diagram of the protocol used for purification of the full-size TatB proteins after overexpression in *E. coli*. The protein peak visible at 23 min in the reversed phase (*RP*)-*HPLC* profile of wild-type TatB shown as example was recovered and analysed by mass spectroscopy (*MALDI-TOF MS*). (**C**) Aliquots from each preparation corresponding to 1-2 μg purified protein were loaded on a 15% SDS-PAA gel and visualised by Coomassie colloidal staining (*lane 1* - wild-type TatB, *lane 2* - antigen, *lane 3* - TatB,E10C).

If such affinity-purified antibodies were added to thylakoid vesicles utilized in *in thylakoido* protein transport experiments, membrane transport of Tat substrates like pOEC16, the precursor of the 16 kDa subunit of the oxygen-evolving system, was abolished (Fig. 2). Supplementation of the assays with TatB obtained from cell-free synthesis with the *Rapid Translation System* (RTS) resulted in partial recovery of thylakoid transport of the Tat substrate demonstrating that protein transport by the thylakoidal Tat pathway can in principle be realised also with soluble TatB that is added externally (Fig. 2). However, the degree of transport recovery was only low in these experiments, in contrast to the quantitative reconstitution observed with TatA in such assays (Hauer et al., 2013). Moreover, the amount of TatB protein obtained by *in vitro* translation varies to some extent and can furthermore not be quantified which limits the significance of the data to a qualitative level.

**Fig. 2.**
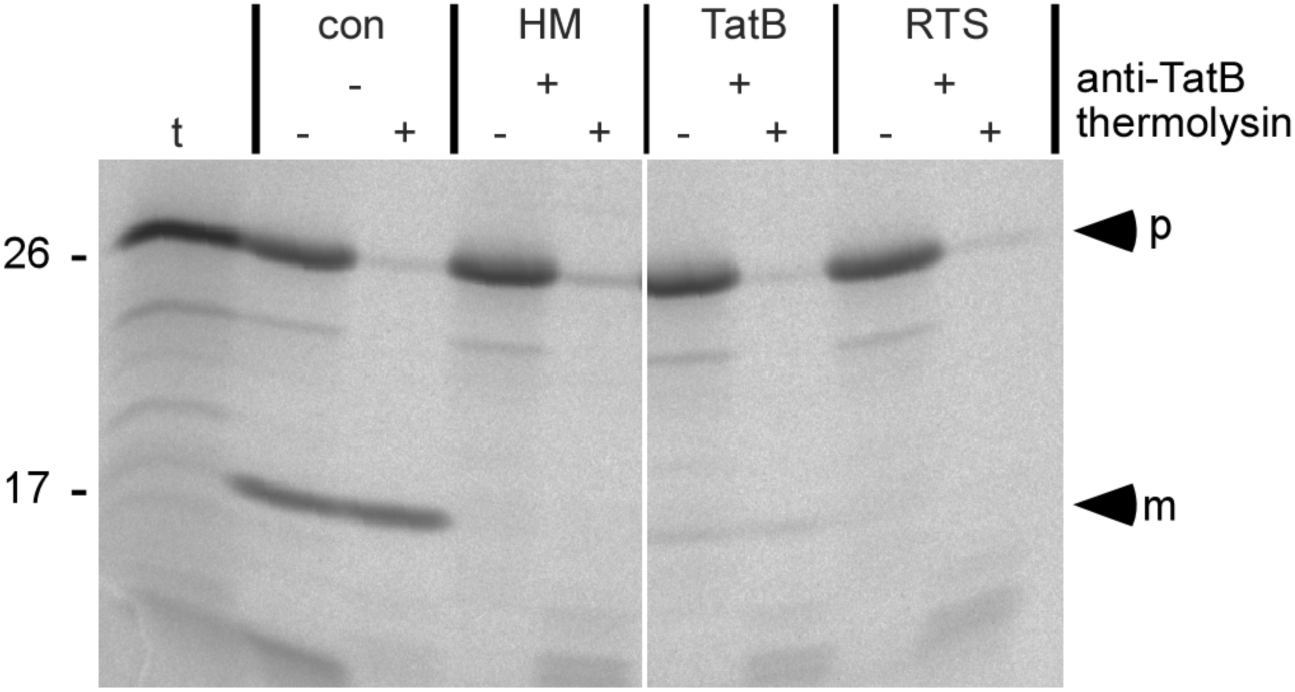
Reconstitution of anti-TatB treated thylakoids with soluble TatB. The precursor protein of OEC16 (*pOEC16*) was synthesized by *in vitro* transcription/ translation of the corresponding cDNA in the presence of [^35^S]-methionine. 5 μl of the translation assay was incubated with isolated pea thylakoids that were either mock-treated (*con*) or treated with affinity-purified antibodies raised against TatB (*HM*). In the reconstitution assays, the anti-TatB treated thylakoids were supplemented with TatB (*TatB*), which was obtained by *in vitro* transcription/translation with the wheat germ rapid translation system RTS, or with RTS containing an empty vector control (*RTS*). After the import reaction for 10 min at 25°C in the light, thylakoids were washed with HM buffer (10 mM Hepes/KOH, pH 8.0; 5 mM MgCl_2_) and either treated with thermolysin (200 μg/ml, 30 min on ice, *lanes +*), or mock treated (*lanes -*). Stoichiometric amounts of each fraction corresponding to 7.5 μg of chlorophyll were separated on a 10-17.5% SDS-PAA gradient gel and visualised by phosphorimaging. In lane *t*, 1 μl of the pOEC16 *in vitro* translation assay was loaded. The positions of the OEC16 precursor (*p*) and mature proteins (*m*) are indicated by *arrowheads*. The mobility of marker proteins (given in kDa) is shown on the left side. The *white space* indicates that in this figure separate parts from the same gel were mounted together.

### Heterologous overexpression and purification of wild-type and mutant TatB

Quantitative analyses require chemical amounts of TatB, which were again obtained by heterologous overexpression in *E. coli* (Fig. 1). In addition to wild-type TatB, a mutant derivative (TatB,E10C) carrying a cysteine residue instead of a glutamate at position 10 of the mature protein was generated in the same manner. Both TatB proteins were synthesized in *E. coli* as fusion polypeptides carrying N-terminal His/S-tags suitable for Ni-affinity chromatography (Fig. 1A). After chromatography the tags were removed by BrCN cleavage and the proteins were further purified by preparative SDS-PAGE and RP-HPLC (Fig. 1B). The quality of the purified proteins, which were routinely obtained at concentrations of about 2 μM in detergent-free buffer, was controlled by SDS-PAGE and mass spectrometry (e.g., Fig. 1 B,C).

### Reconstitution of thylakoidal Tat transport by purified soluble TatB

Both TatB proteins as well as the purified antigen were then subjected to *in thylakoido* reconstitution assays performed with antiTatB-treated thylakoids using again radiolabelled pOEC16 as transport substrate. While in the buffer control membrane transport of pOEC16 into the antibody-treated thylakoid vesicles was abolished, supplementation of the assays with 0.5 - 1.5 μM wild-type TatB led to the recovery of thylakoidal Tat transport (Fig. 3A). Membrane transport of the Tat substrate was considerably stronger in this case than in the reconstitution assays employing TatB obtained by *in vitro* translation (Fig. 2) demonstrating that wild-type TatB can be purified in catalytically active form that is principally able to replace intrinsic TatB in function. However, transport was still less efficient than with mock-treated control thylakoids which might be indicative of a time-consuming assembly step for the externally added protein into the receptor complex. We have therefore examined if the transport activity in the reconstitution assays depends on the time of preincubation of anti-TatB-treated thylakoids with purified TatB prior to the actual transport experiment. Although some minor variation could be observed with time periods ranging from 0 - 20 min, there was no clear-cut correlation of preincubation time and degree of transport reconstitution (Fig. S1) suggesting that the time period of preincubation does not have a major impact on Tat transport recovery. Yet, to be on the safe side we maintained a preincubation time of 5 min in all our experiments.

**Fig. 3.**
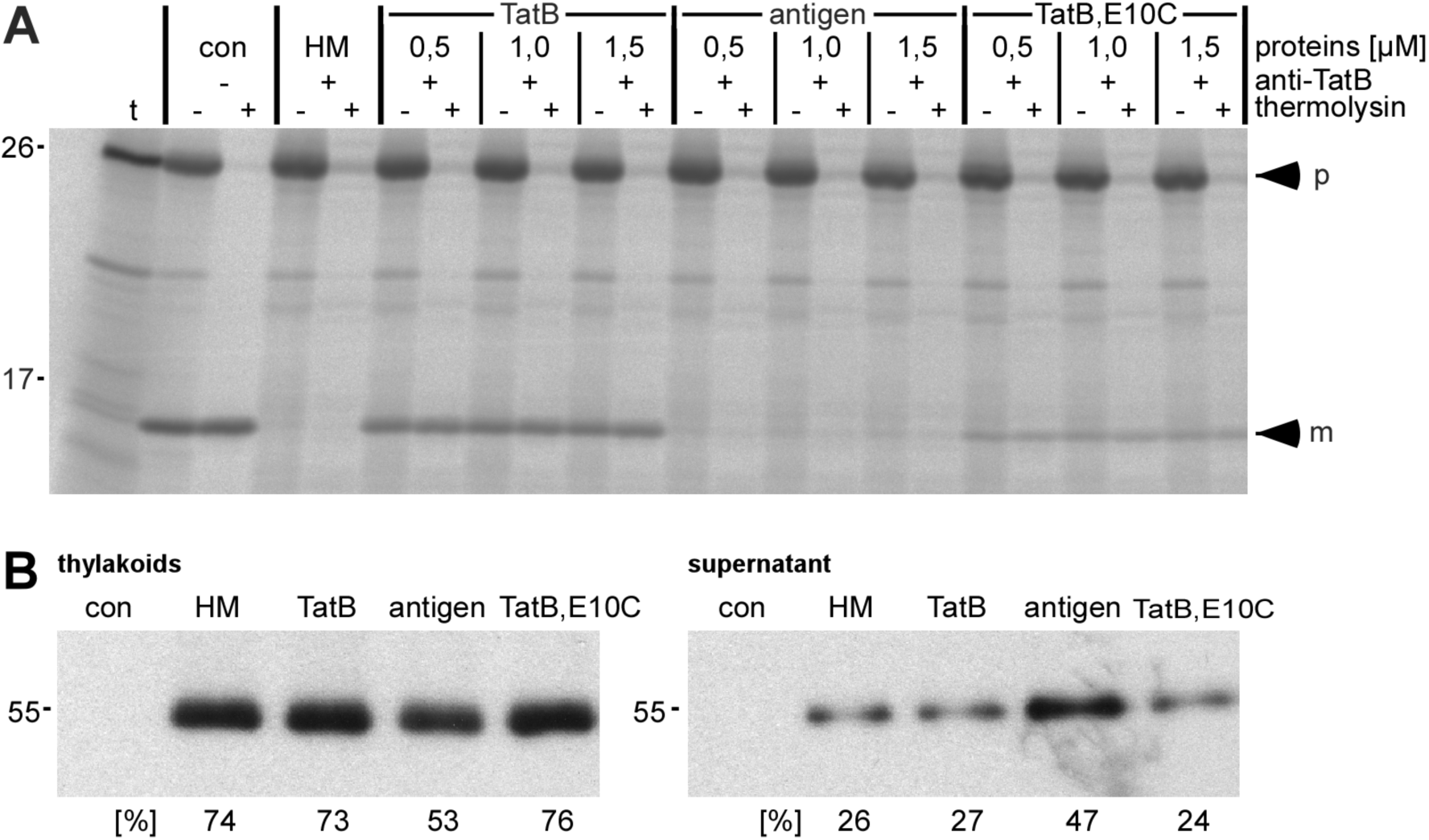
Persistence of antibody inhibition. (**A**) *In thylakoido* assays containing anti-TatB treated thylakoids were supplemented with either HM buffer (*HM*) or with 0.5 μM, 1 μM, and 1.5 μM of either purified TatB (*TatB*), TatB antigen (*antigen*), or mutant *TatB,E10C*. After 5 min preincubation on ice the transport reaction was started by adding radiolabelled pOEC16 and analysed as described in the legend to Fig. 2. (**B**) Transport assays analogous to those shown in (**A**) were centrifuged for 4 min at 10.000 x g. Stoichiometric amounts of the pellet (*thylakoids*) and *supernatant* fractions corresponding to 7.5 µg chlorophyll each were separated by 15% SDS-PAGE and transferred to PVDF membranes. To determine the amount of the anti-TatB antibodies in the two fractions the membranes were incubated for 1 h with horseradish peroxidase-coupled anti-rabbit IGG and subjected to ECL (*enhanced chemiluminescence*) detection. The relative amounts of the heavy chains of anti-TatB antibodies in the two fractions were separately calculated for each reconstitution assay and are given in terms of percentage below each lane.

Reconstitution experiments performed with 0.5 - 1.5 μM purified TatB antigen, which lacks the N-terminal TMH, did not lead to any notable membrane transport of pOEC16 (Fig. 3A). This suggests already that the transport activity observed after supplementing the assays with full-size TatB was caused by the activity of the protein added and not the result of competitive release of the antibodies from intrinsic TatB. This conclusion could further be confirmed by determining the extent of antibody release by the externally added proteins. The antigen was by far more efficient in liberating the antibodies from intrinsic TatB than either wild-type TatB or TatB,E10C (Fig. 3B). While in the presence of either of the two full-size TatB proteins the amount of anti-TatB antibodies liberated was in the same low range as in the buffer control leaving the majority of the antibodies bound to the thylakoids, supplementation of the assays with the antigen at the same concentration led to approximately 50% antibody release into the supernatant (Fig. 3B). Yet, the Tat transport activity of such antigen-treated thylakoids was negligible (Fig. 3A) which finally rules out that the transport recovery observed in the presence of full-size TatB was caused by reactivation of the intrinsic TatB acitivity.

Remarkably, even mutant TatB,E10C was able to reconstitute thylakoid transport of pOEC16 (Fig. 3A). Transport recovery was significantly lower in this case than in the presence of wild-type TatB but clearly above the background activity observed in the presence of antigen. This result was entirely unexpected because according to literature data (Ma et al., 2018) the glutamate residue at position 10 within TMH (termed E11 in there) is essential for the assembly of TatB into thylakoidal TatBC receptor complexes. Such assembly is generally assumed to be a prerequisite for TatB function which in turn suggests that assembly of the mutant protein is not totally abolished but probably takes place only at a level below the detection limit of the published assay. How such poor assembly can account for the low but considerable transport activity observed here remains enigmatic so far. Yet, it suggests that the reduced transport of pOEC16 in the reconstitution assays comprising TatB,E10C might reflect the low assembly competence of the mutant TatB protein rather than being the result of lower catalytic activity.

### Quantification of TatB demand

As the first step to quantify the demand of TatB during membrane transport of pOEC16, the appropriate incubation time had to be determined. For this purpose, we have compared the time-course of thylakoid transport of the Tat substrate pOEC16 in the reconstituted system with that of mock-treated control thylakoids. As expected, membrane transport in the control reaction was very fast showing saturation of the accumulation of mature OEC16 in the thylakoid lumen already after 15 min (Fig. 4). In contrast, in the reconstitution assays a slow but permanent increase of the amount of mature OEC16 could be observed which did not reach a saturation level within the time of analysis (20 min). To allow for sufficient signal strength on the one hand but still allowing comparison with mock-treated control thylakoids on the other hand, we have chosen an incubation time of 10 min for the subsequent experiments.

**Fig. 4.**
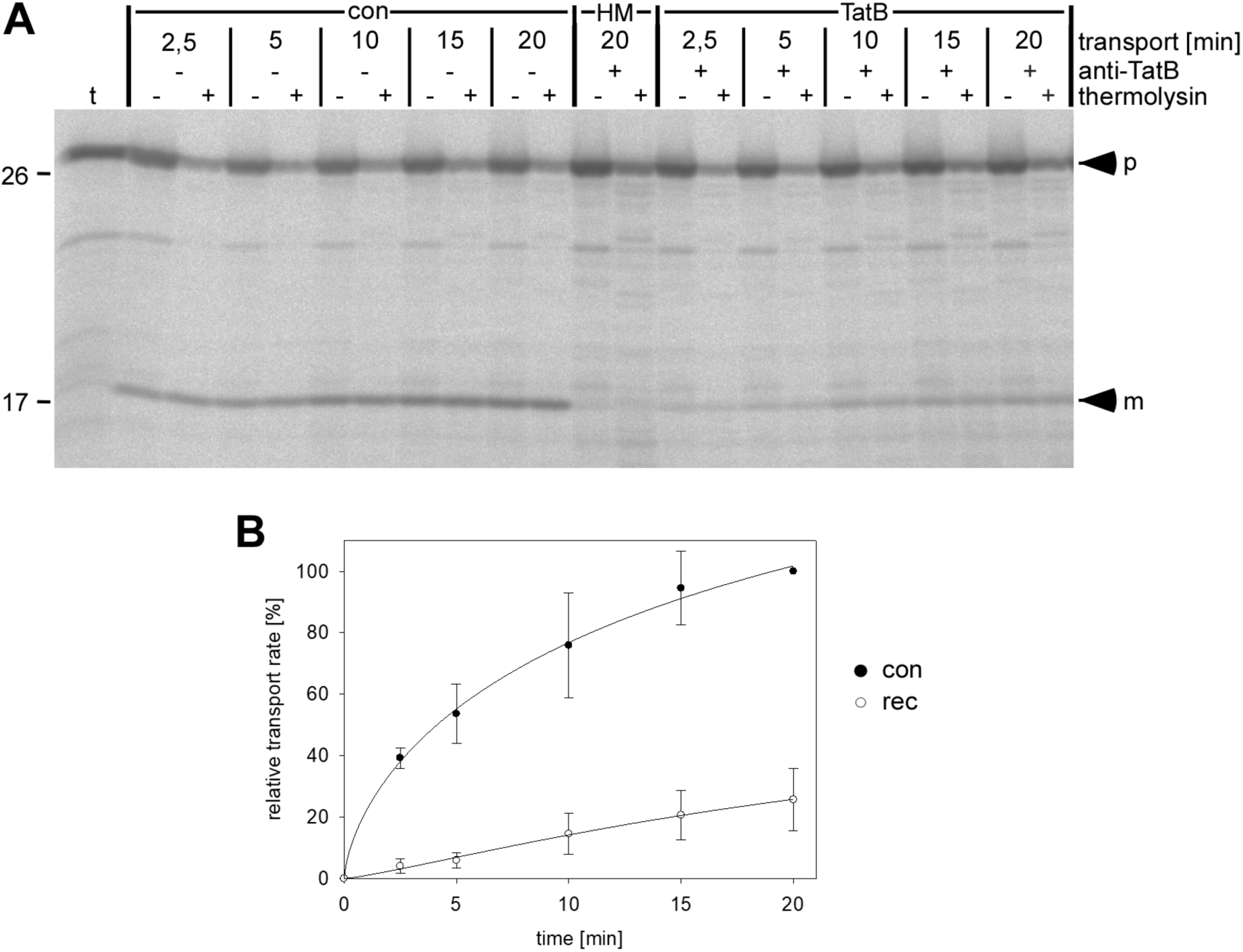
Time-course of Tat transport after TatB reconstitution. (**A**) Mock-treated thylakoids (*con*) or anti-TatB treated thylakoids supplemented with either HM buffer (*HM*) or with 0.5 μM purified *TatB* were incubated with radiolabeled pOEC16 at 25°C in the light for the time periods indicated on top of the lanes. All further processing steps were performed as described in the legend to Fig. 2. (**B**) Quantification of the amount of translocated OEC16 from repeated reconstitution experiments identical to those shown in (**A**). The results obtained with mock-treated thylakoids are given with *filled circles*, those of the reconstitution assays with *open circles*. Both mean values and standard deviations were calculated from three (n = 3) independently performed experiments. For each time-point, the relative amount of mature OEC16 (*m*) was calculated in terms of percentage of the value obtained in the 20 min control reaction.

The quantitative reconstitution experiments were then carried out by supplementing antibody-treated thylakoids with soluble wild-type TatB at concentrations ranging from 0.025 μM to 1.5 μM (Fig. 5A). For each concentration three independent experiments were conducted. The accumulation of mature OEC16 in the thylakoid lumen was quantified and related to the amount obtained in control experiments with mock-treated thylakoids performed in parallel. Plotting these “relative transport rates” against the respective concentration of soluble TatB in the assays yielded a sigmoidal curve that is typical for enzymes exhibiting cooperative activity. While at low TatB concentration (≤ 0.075 μM) transport could hardly be detected, any higher concentration of TatB in the assays led to strongly increased Tat transport activity until at approximately 0.5 μM TatB maximal transport was achieved (Fig. 5B). Under these conditions, the transport rate reached 20 - 25% of that obtained with mock-treated control thylakoids.

**Fig. 5.**
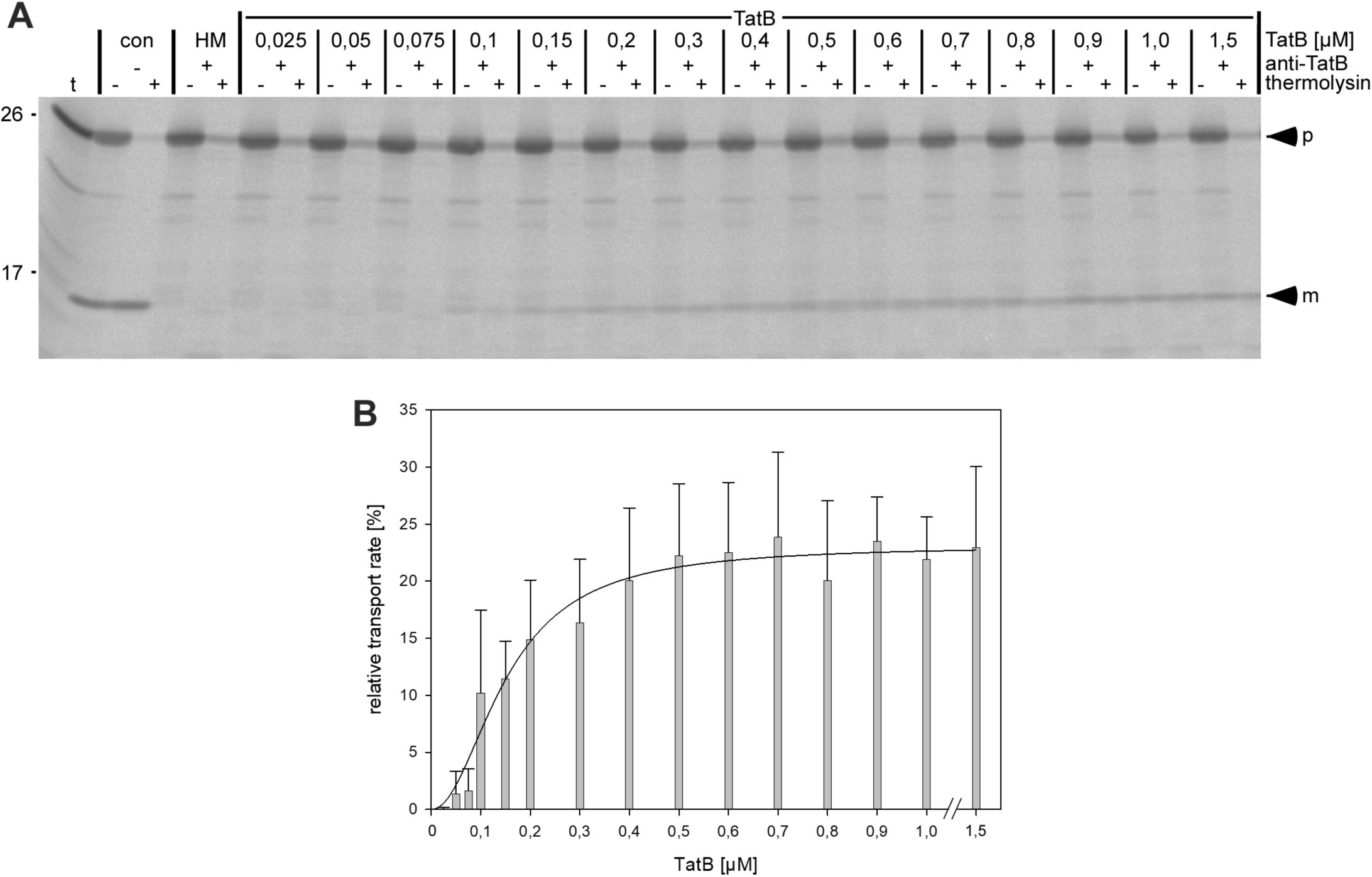
Quantification of TatB requirement. (**A**) *In thylakoido* assays containing anti-TatB treated thylakoids were supplemented with purified soluble TatB at the concentrations indicated on the top of the lanes and incubated with radiolabeled pOEC16 for 10 min at 25°C in the light. (**B**) Quantification of the amount of translocated protein from repeated transport reactions identical to those shown in (**A**). For each TatB concentration, the relative amount of translocated protein (*m*) was calculated in terms of percentage of the corresponding control reaction with mock-treated thylakoids (*con*). Both mean values and standard deviations were calculated from three (n = 3) independently repeated experiments. The fit corresponds to a sigmoidal function according to the Hill equation v = (V_max_ * [TatB]^n^) / (Km ^n^ + [TatB] ^n^). For further details, see the legends to Figs. 2 and 4.

## Discussion

It was the goal of this study to establish an experimental system for the functional analysis of thylakoidal TatB, a component of the TatBC receptor structure of the thylakoidal Twin-arginine protein translocation pathway. Using an approach that had been established earlier for the analysis of thylakoidal TatA, namely antibody inhibition of the intrinsic Tat activity followed by transport reconstitution by supplementing the assays with the respective Tat component, membrane transport of the authentic Tat substrate pOEC16 could be reconstituted to a considerable degree demonstrating that purified soluble TatB is catalytically active and principally able to replace intrinsic TatB in function.

### Conditions for functional TatB reconstitution

Such transport reconstitution could not necessarily be expected given the structural prerequisites for TatB function. The protein is part of the membrane-integral, heterooligomeric TatBC receptor complex of approximately 560 - 700 kDa (Berghöfer and Klösgen, 1999; Cline and Mori, 2001) which apparently comprises TatB and TatC in equimolar amounts (Bolhuis et al., 2001; Jack et al., 2001; Jakob et al., 2009). Functional reconstitution therefore depends not only on the correct insertion of the externally added soluble TatB protein into the membrane but presumably also on its assembly into such membrane complexes. Hence, it is probably not surprising that Tat transport in the reconstitution assays reaches only 20 - 25% of the maximal rate obtained with mock-treated thylakoid vesicles (Figs. 4 and 5). Quantitative reconstitution, as observed for TatA in similar experiments (Hauer et al., 2013), might have been expected in first place considering the structural similarity of TatA and TatB (Yen et al., 2002). However, for TatA no demand for a comparable assembly with other Tat components has been described.

The requirement of TatB assembly might be the reason also for the low level of reconstitution observed for the TatB derivative TatB,E10C (Fig. 3A). In fact, in this case reconstitution was even unexpected in view of an earlier publication (Ma et al., 2018) demonstrating lack of assembly of this mutant (termed E11C) into TatBC complexes. However, our data suggest that assembly of the TatB,E10C mutant might not be completely abolished but to take place at levels that are below the detection limit of the Blue Native-PAGE analysis published by Ma et al. (2018). In line with that, the corresponding mutant of *E. coli*, TatB,E8A, likewise showed deviating transport properties in independent analyses (Hicks et al., 2003; Alcock et al., 2016).

In any case, TatB assembly demands for the availability of potential binding sites and maybe even an assembly machinery. It is yet unknown if assembly of the externally added TatB protein takes place into already existing TatBC complexes, e.g., by replacing antibody-blocked TatB subunits or by occupying free binding sites of partially assembled complexes. Alternatively, such complexes might also be assembled *de novo* making use of potentially surplus TatC molecules in the membrane. In this case, the availability of TatC would be a limiting factor, i.e., the degree of transport recovery would depend on the availability of sufficient amounts of free TatC in the membrane. It might therefore be worthwhile trying to increase the transport efficiency in the reconstituted system by supplementing the assays additionally with TatC. Unfortunately though, the extraordinary hydrophobic nature of this protein complicates its purification and affects its solubility in detergent-free buffers which prevented such experiments to date.

### TatB demand during thylakoidal protein transport

Quantification of the amount of TatB required to facilitate thylakoid transport of the authentic Tat substrate pOEC16 yielded a rather unexpected result. Although the only moderate levels of TatB reconstitution demand for particularly careful interpretation of the data, the overall similarity to the results obtained with TatA in such experiments (Hauer et al., 2013) is still remarkable. Both Tat subunits show a sigmoidal activity curve indicative of cooperative catalytic activity (Fig. 5B) and even the Km-value calculated from these data for TatB (approximately 0.15 μM) is remarkably similar to that determined for TatA (Km = 0.12 μM; Hauer et al. 2013). Thus, both Tat subunits are apparently required in comparable concentration to achieve 50% transport recovery in such reconstitution assays suggesting that TatA and TatB might be present also in equimolar amounts in the thylakoid membrane. This result stands in clear contrast to the excess amount of TatA over TatB and TatC (≥ 20:1) postulated in numerous publications (e.g., Jack et al., 2001 Sargent et al., 2001; Berks et al., 2003; Celedon and Cline, 2012). However, while in *E. coli* an excess of TatA is generally assumed (Jack et al., 2001), the stoichiometry of the Tat subunits in plants is still disputed. Both substoichiometric (Jakob et al., 2009), stoichiometric (Mori et al., 2001; Jakob et al., 2009), as well as excess amounts of TatA (Celedon and Cline, 2012) compared with TatB and TatC were described depending on the method used for analysis and/or the plant species studied. It should be noted though that in most instances steady-state levels of proteins had been determined without considering their actual activity status. In contrast, in the experiments presented here as well as in Hauer et al. (2013), solely the demand of catalytically active TatA and TatB was determined, although we have no proof, of course, that our protein preparations are entirely free from inactive polypeptides. Still, the observation of presumably comparable demands for TatA and TatB upon Tat transport challenges some of the current working models.

## Materials and methods

### Cloning of TatB, TatB,E10C, and TatB antigen

The cDNA fragment encoding mature TatB from *Pisum sativum* (gene accession number AF284760) was amplified by PCR using a cDNA library generated from seedling leaves as template and the following primers: 5’ ttt cca tgg cgt ctc tct ttg ggg ttg 3’ and 5’ ttt ctc gag cta taa atc cga agg taa cg 3’. The amplified fragment was cut with NcoI and XhoI and cloned with two different vectors (linearized with NcoI and XhoI), (i) the pIVEX 1.3 WG vector suitable for cell-free synthesis of the protein with the *Rapid Translation System* (5PRIME, Hamburg, Germany) and (ii) the bacterial expression vector pET-30a (Novagen). The latter yielded a chimeric TatB construct carrying a combined His-/S-peptide-tag ending with a BrCN cleavage site immediately upstream of the N-terminus of mature TatB (Fig. 1). To obtain TatB,E10C QuikChange® Site-Directed Mutagenesis (Stratagene, La Jolla, CA, USA) was performed using the primers 5’ ctc tct ttg ggg ttg gag cac ctt gtg ctt tgg taa ttg ggg ttg tgg c 3’ and 5’ gcc aca acc cca att acc aaa gca caa ggt gct cca acc cca aa gaga g 3’. The fragment encoding the TatB antigen comprising all but the TMH of mature TatB was amplified with the primers 5’ ttt cca tgg gtc cta aag gtc ttg ctg 3’ and 5’ ttt ctc gag taa atc cga agg taa cga cg 3’ and transferred into pET28 using the restriction sites NcoI and XhoI for cloning.

### Heterologous overexpression and purification of TatB, TatB,E10C, and TatB antigen

The N-terminally His-/S-peptide-tagged TatB and TatB,E10C, as well as the C-terminally His-tagged antigen were obtained by overexpression in *E. coli* strain BL21 (DE3) over night at 37°C in M9 minimal medium. The bacterial suspension was supplemented with 5 volumes binding buffer (20 mM Hepes, 500 mM NaCl, 20 mM Imidazole, 6 M GuanidinHCl, pH 7.5) and incubated for 60 min at 4°C. After ultracentrifugation for 60 min at 140.000 g, the supernatant was applied to Ni^2+^-affinity chromatography (HiTrap Chelating HP columns, GE Healthcare) according to the manufacturer’s instructions. All TatB variants were recovered with four column volumes elution buffer each (20 mM Hepes, 500 mM NaCl, 500 mM Imidazole, 6 M GuanidinHCl, pH 7.5). Removal of the N-terminal His/S-tags from the full-size TatB proteins by BrCN cleavage, preparative SDS-PAGE, Zinc-imidazole staining, electroelution, and reversed phase HPLC followed the protocol described in Hauer et al. (2013). The HPLC fractions containing TatB were combined and dialysed at 4°C over night against HM buffer (10 mM Hepes-KOH, pH 8.0; 5 mM MgCl_2_). After removal of aggregates by ultracentrifugation (20 min, 160.000 x g, 4°C) the protein solutions were stored at 4°C.

### In thylakoido reconstitution assays

Isolation of chloroplasts and thylakoids from pea seedlings (*Pisum sativum* var. Feltham First) followed the protocol described by Hou et al. (2006). *In thylakoido* protein transport experiments with radiolabelled precursor proteins were carried out according to Marques et al. (2003) with the following modifications: to inhibit protein transport by the Tat pathway, thylakoids were incubated for 45 min on ice with affinity-purified antibodies against TatB, washed twice with HM-buffer, and resuspended in HM buffer supplemented with TatB, TatB,E10C, or antigen. After preincubation for 5 min on ice, the actual import reaction was started by the addition of the radiolabelled Tat substrate pOEC16 obtained by *in vitro* translation. Subsequent processing of the assays was performed as described (Marques et al., 2003).

### Miscellaneous

Gel electrophoresis of proteins under denaturing conditions was carried out according to Laemmli et al. (1970). The gels were exposed to phosphorimager screens and analysed with the Fujifilm FLA-3000 (Fujifilm, Düsseldorf, Germany) using the software packages BASReader (version 3.14) and AIDA (version 3.25; Raytest, Straubenhardt, Germany). Cell-free synthesis of proteins was carried out with the *Rapid Translation System (RTS) 100 wheat germ CECF kit* according to the manufacturer’s instructions (5PRIME, Hamburg, Germany). Protein concentration was determined according to Bradford (1976). Western analysis was performed as described by Jakob et al. (2009). All other methods followed published protocols (Sambrook et al., 2001).

## Acknowledgements

We thank Norbert Ruthenberg for his help in the preparation of the TatB antigen and Birgit Kretschmann for excellent technical assistance.

---

**Fig. S1.**
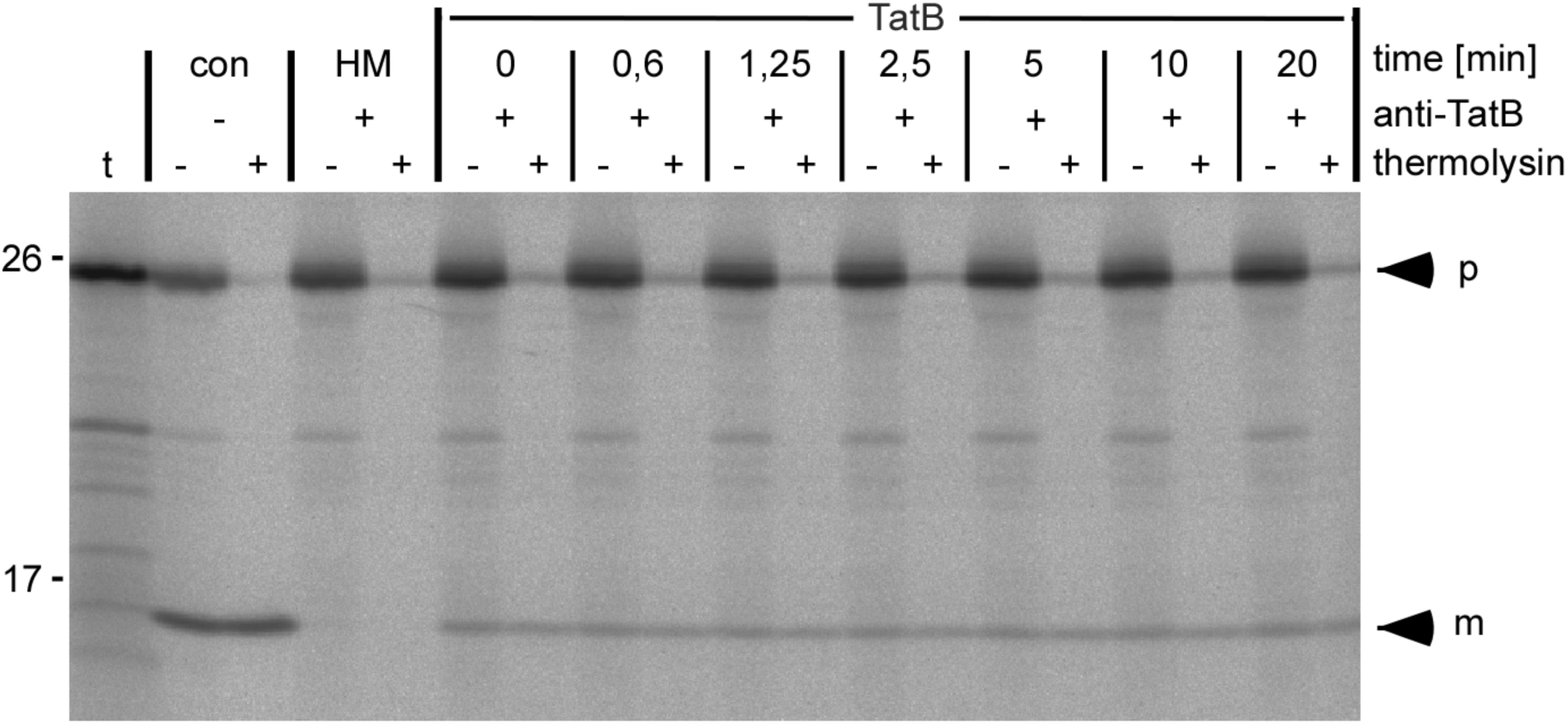
*Impact of TatB preincubation on transport reconstitution In thylakoido* assays containing antiTatB-treated thylakoids were supplemented with 0.5 μM purified soluble TatB and incubated on ice for the time periods indicated on top of the lanes. The import reaction was started by supplementing the assays with radiolabelled pOEC16 and subsequently analysed as described in the legend to Fig. 2.

## Footnotes

1 The abbreviations used are: Tat, twin-arginine translocation; OEC16, 16 kDa subunit of the oxygen-evolving system associated with photosystem II; RTS, Rapid Translation System; Hepes, 2-[4-(2-hydroxyethyl)piperazin-1-yl]ethanesulfonic acid; BrCN, cyanogen bromide

## References

F. Alcock, P.J. Stansfeld, H. Basit, J. Habersetzer, M.A.B. Baker, T. Palmer, M.I. Wallace, B.C. Berks, Assembling the Tat protein translocase, (2016) 1–28. doi:10.7554/eLife.20718.

J. Berghöfer, R.B. Klösgen, Two distinct translocation intermediates can be distinguished during protein transport by the TAT (Δph) pathway across the thylakoid membrane, FEBS Lett. 460 (1999) 328–332. doi:10.1016/S0014-5793(99)01365-4.

B.C. Berks, T. Palmer, F. Sargent, The Tat protein translocation pathway and its role in microbial physiology, Adv. Microb. Physiol. 47 (2003) 187–254. doi:10.1016/S0065-2911(03)47004-5.

A. Bolhuis, J.E. Mathers, J.D. Thomas, C.M. Barrett, C. Robinson, TatB and TatC form a functional and structural unit of the twin-arginine translocase from Escherichia coli., J. Biol. Chem. 276 (2001) 20213–9. doi:10.1074/jbc.M100682200.

J.M. Celedon, K. Cline, Stoichiometry for binding and transport by the twin arginine translocation system., J. Cell Biol. 197 (2012) 523–34. doi:10.1083/jcb.201201096.

S.A. Clark, S.M. Theg, A folded protein can be transported across the chloroplast envelope and thylakoid membranes., Mol. Biol. Cell. 8 (1997) 923–34. http://www.ncbi.nlm.nih.gov/pubmed/9168475 (accessed June 24, 2018).

K. Cline, Mechanistic Aspects of Folded Protein Transport by the Twin Arginine Translocase (Tat), J. Biol. Chem. 290 (2015) 16530–16538. doi:10.1074/jbc.R114.626820.

K. Cline, M. McCaffery, Evidence for a dynamic and transient pathway through the TAT protein transport machinery., EMBO J. 26 (2007) 3039–3049. doi:10.1038/sj.emboj.7601759.

K. Cline, W.F. Ettinger, S.M. Theg, Protein-specific energy requirements for protein transport across or into thylakoid membranes. Two lumenal proteins are transported in the absence of ATP, J. Biol. Chem. 267 (1992) 2688–2696.

K. Cline, H. Mori, ThylakoidΔpH-dependent precursor proteins bind to a cpTatC-Hcf106 complex before Tha4-dependent transport, J. Cell Biol. 154 (2001) 719–729. doi:10.1083/jcb.200105149.

C. Dabney-Smith, H. Mori, K. Cline, Requirement of a Tha4-conserved Transmembrane Glutamate in Thylakoid Tat Translocase Assembly Revealed by Biochemical Complementation, J. Biol. Chem. 278 (2003) 43027–43033. doi:10.1074/jbc.M307923200.

E. Fan, M. Jakob, R.B. Klösgen, One signal is enough: Stepwise transport of two distinct passenger proteins by the Tat pathway across the thylakoid membrane, Biochem. Biophys. Res. Commun. 398 (2010) 438–443. doi:10.1016/j.bbrc.2010.06.095.

S. Frielingsdorf, M. Jakob, R.B. Klösgen, A stromal pool of TatA promotes Tat-dependent protein transport across the thylakoid membrane., J. Biol. Chem. 283 (2008) 33838–45. doi:10.1074/jbc.M806334200.

U. Gohlke, L. Pullan, C.A. McDevitt, I. Porcelli, E. de Leeuw, T. Palmer, H.R. Saibil, B.C. Berks, The TatA component of the twin-arginine protein transport system forms channel complexes of variable diameter., Proc. Natl. Acad. Sci. U. S. A. 102 (2005) 10482–6. doi:10.1073/pnas.0503558102.

J. Habersetzer, K. Moore, J. Cherry, G. Buchanan, P.J. Stansfeld, T. Palmer, J. Cherry, G. Buchanan, S. Pj, T. Palmer, T. Palmer, Substrate-triggered position switching of TatA and TatB during Tat transport in Escherichia coli, (2017).

R.S. Hauer, R. Freudl, J. Dittmar, M. Jakob, R.B. Klösgen, How to achieve Tat transport with alien TatA, Sci. Rep. 7 (2017) 1–13. doi:10.1038/s41598-017-08818-w.

R.S. Hauer, R. Schlesier, K. Heilmann, J. Dittmar, M. Jakob, R.B. Klösgen, Enough is enough: TatA demand during Tat-dependent protein transport., Biochim. Biophys. Acta. 1833 (2013) 957–65. doi: 10.1016/j.bbamcr.2013.01.030.

M.G. Hicks, E. De Leeuw, I. Porcelli, G. Buchanan, B.C. Berks, T. Palmer, The Escherichia coli twin-arginine translocase: Conserved residues of TatA and TatB family components involved in protein transport, FEBS Lett. 539 (2003) 61–67. doi:10.1016/S0014-5793(03)00198-4.

B. Hou, E.S. Heidrich, D. Mehner-Breitfeld, T. Brüser, The TatA component of the twin-arginine translocation system locally weakens the cytoplasmic membrane of Escherichia coli upon protein substrate binding, J. Biol. Chem. (2018). doi:10.1074/jbc.RA118.002205.

Y. Hu, E. Zhao, H. Li, B. Xia, C. Jin, Solution NMR structure of the TatA component of the twin-arginine protein transport system from gram-positive bacterium bacillus subtilis, J. Am. Chem. Soc. 132 (2010) 15942–15944. doi:10.1021/ja1053785.

P.J. Hynds, D. Robinson, C. Robinson, The Sec-independent twin-arginine translocation system can transport both tightly folded and malfolded proteins across the thylakoid membrane, J. Biol. Chem. 273 (1998) 34868–34874. doi:10.1074/jbc.273.52.34868.

R.L. Jack, F. Sargent, B.C. Berks, G. Sawers, T. Palmer, Constitutive expression of Escherichia coli tat genes indicates an important role for the twin-arginine translocase during aerobic and anaerobic growth., J. Bacteriol. 183 (2001) 1801–4. doi:10.1128/JB.183.5.1801-1804.2001.

M. Jakob, S. Kaiser, M. Gutensohn, P. Hanner, R.B. Klösgen, Tat subunit stoichiometry in Arabidopsis thaliana challenges the proposed function of TatA as the translocation pore., Biochim. Biophys. Acta. 1793 (2009) 388–94. doi: 10.1016/j.bbamcr.2008.09.006.

R.B. Klösgen, I.W. Brock, R.G. Herrmann, C. Robinson, Proton gradient-driven import of the 16 kDa oxygen-evolving complex protein as the full precursor protein by isolated thylakoids, Plant Mol. Biol. 18 (1992) 1031–1034. doi:10.1007/BF00019226.

Q. Ma, K. Fite, C.P. New, C. Dabney-Smith, Thylakoid-integrated recombinant Hcf106 participates in the chloroplast twin arginine transport system, Plant Direct. 2 (2018). doi:10.1002/pld3.90.

J.P. Marques, I. Dudeck, R.B. Klösgen, Targeting of EGFP chimeras within chloroplasts., Mol. Genet. Genomics. 269 (2003) 381–7. doi:10.1007/s00438-003-0846-y.

H. Mori, K. Cline, A twin arginine signal peptide and the pH gradient trigger reversible assembly of the thylakoidΔpH/Tat translocase, J. Cell Biol. 157 (2002) 205–210. doi:10.1083/jcb.200202048.

M. Müller, R.B. Klösgen, The Tat pathway in bacteria and chloroplasts (review), Mol. Membr. Biol. 22 (2005) 113–121.

P. Natale, T. Brüser, A.J.M. Driessen, Sec- and Tat-mediated protein secretion across the bacterial cytoplasmic membrane — Distinct translocases and mechanisms, 1778 (2008) 1735–1756. doi: 10.1016/j.bbamem.2007.07.015.

P. Pettersson, W. Ye, M. Jakob, F. Tannert, R.B. Klösgen, L. Mäler, Structure and dynamics of plant TatA in micelles and lipid bilayers studied by solution NMR, FEBS J. 285 (2018) 1886–1906. doi:10.1111/febs.14452.

S. Ramasamy, R. Abrol, C.J.M. Suloway, W.M. Clemons, The glove-like structure of the conserved membrane protein tatc provides insight into signal sequence recognition in twin-arginine translocation, Structure. 21 (2013) 777–788. doi:10.1016/j.str.2013.03.004.

F. Rodriguez, S.L. Rouse, C.E. Tait, J. Harmer, A. De Riso, C.R. Timmel, M.S.P. Sansom, B.C. Berks, J.R. Schnell, Structural model for the protein-translocating element of the twin-arginine transport system., Proc. Natl. Acad. Sci. U. S. A. 110 (2013) E1092–101. doi:10.1073/pnas.1219486110.

S.E. Rollauer, M.J. Tarry, J.E. Graham, M. Jääskeläinen, F. Jäger, S. Johnson, M. Krehenbrink, S.-M. Liu, M.J. Lukey, J. Marcoux, M.A. McDowell, F. Rodriguez, P. Roversi, P.J. Stansfeld, C. V Robinson, M.S.P. Sansom, T. Palmer, M. Högbom, B.C. Berks, S.M. Lea, Structure of the TatC core of the twin-arginine protein transport system., Nature. 492 (2012) 210–4. doi:10.1038/nature11683.

J. Sambrook, D.W. Russell, Molecular Cloning: A Laboratory Manual, 3rd ed. Cold Spring Harbor Laboratories, Cold Spring Harbor, NY, 2001.

F. Sargent, U. Gohlke, E. De Leeuw, N.R. Stanley, T. Palmer, H.R. Saibil, B.C. Berks (2001) Purified components of the Escherichia coli Tat protein transport system form a double-layered ring structure. Eur J Biochem 268(12):3361–3367

M.R. Yen, Y.H. Tseng, E.H. Nguyen, L.F. Wu, M.H. Saier, Sequence and phylogenetic analyses of the twin-arginine targeting (Tat) protein export system, Arch. Microbiol. 177 (2002) 441–450. doi:10.1007/s00203-002-0408-4.

Y. Zhang, L. Wang, Y. Hu, C. Jin, Solution structure of the TatB component of the twin-arginine translocation system., Biochim. Biophys. Acta. 1838 (2014) 1881–8. doi: 10.1016/j.bbamem.2014.03.015.

